# Sialidase-Mediated Desialylation Regulating EGFR Phosphorylation and Signal Flux

**DOI:** 10.64898/2026.05.26.728000

**Authors:** Hongyi Liu, Effram Wei, Ding Chiao Lin, Hui Zhang

**Affiliations:** Department of Pathology, Johns Hopkins University School of Medicine, Baltimore, MD 21231, USA; Department of Chemical and Biomolecular Engineering, Johns Hopkins University Whiting School of Engineering, Baltimore, MD 21218, USA

**Keywords:** glycosylation, phosphorylation, proteomic, Sialidase, EGFR, ccRCC

## Abstract

Protein glycosylation and phosphorylation are fundamental post-translational modifications (PTMs) that coordinate cellular signaling. While receptor tyrosine kinases (RTKs) like the epidermal growth factor receptor (EGFR) are heavily glycosylated, the systems-level crosstalk between extracellular sialylation and intracellular phosphorylation dynamics remains poorly understood. We employed an integrated TMT18-labeled multi-omics pipeline to simultaneously profile the global proteome, phosphoproteome, and N-glycoproteome of A498 cells. Using enzymatic in-situ remodeling with sialidase, we investigated the signaling response to EGF stimulation and the synergistic effects of desialylation with the tyrosine kinase inhibitor (TKI) gefitinib. Our analysis revealed that cell surface desialylation significantly attenuates EGF-induced signaling, specifically suppressing over 200 phosphosites within the MAPK cascade and actin cytoskeleton organization modules. Comparative profiling demonstrated that sialidase treatment exerts a distinct regulatory program that is non-redundant with canonical TKI inhibition. Stoichiometric analysis confirmed that the depletion of sialylated N-glycoforms at specific EGFR residues (N413, N444) directly correlates with reduced phosphorylation at key activation sites (Y1197). Finally, an integrated glyco-phospho network analysis identified CD44, MET, and integrin signaling hubs as central nodes regulated by the sialylation. This study establishes cell surface sialylation as a critical rheostat for EGFR-mediated signaling flux. By bridging the gap between the extracellular glycoprotein and intracellular kinase networks, we identify glycan remodeling as a potent strategy to sensitize RTK-driven malignancies to therapy. Our findings provide a robust data foundation for developing glycoconjugate-targeted interventions in ccRCC and beyond.

## INTRODUCTION

Protein glycosylation and phosphorylation are two of the most prevalent and functionally significant post-translational modifications (PTMs), orchestrating a vast array of cellular processes including signal transduction, metabolic flux, and cell-cell communication^1,2^. While phosphorylation acts as a rapid molecular switch that propagates intracellular signals through kinase cascades^3,4^, N-linked glycosylation fundamentally shapes the biophysical properties, stability, and interactome of cell-surface receptors^5,6^. Despite their co-occurrence on the majority of signaling hubs^7,8^, the functional crosstalk between the extracellular glycosylation and intracellular phospho-signaling remains one of the most complex yet underexplored frontiers in cell biology.

The clinical significance of this PTM interplay is particularly evident in Clear Cell Renal Cell Carcinoma (ccRCC), the most common and aggressive histological subtype of kidney cancer^9^. ccRCC is characterized by a high degree of metabolic reprogramming and a dependency on growth factor signaling pathways, most notably the epidermal growth factor receptor (EGFR) axes^10–13^. Despite ccRCC dependency on EGFR signaling, EGFR-targeted tyrosine kinase inhibitors (TKIs) have proven clinically ineffective, suggesting unresolved regulatory complexities^14,15^. Genomic studies have long identified the hyperactivation of these pathways as drivers of ccRCC progression^16^. However, genomic data alone fails to capture the dynamic regulatory layers provided by PTMs.

The advent of high-resolution, mass spectrometry (MS)-based proteomics has revolutionized our ability to dissect these complex networks^17^. Multi-omics workflows now allow for the simultaneous quantification of global protein abundance, site-specific phosphorylation, and intact glycopeptides from a single sample^18^. Such system-level approaches are essential for understanding how a specific glycan modification such as the addition of terminal sialic acids can act as a distal regulator of proximal kinase activity. Recent breakthroughs in clinical proteomics and glycoproteomics have revealed that the ccRCC landscape is defined by massive alterations in the glycoproteome^19–23^, where glycosylation of membrane receptors correlates with cancer progression in different cancer types^24,25^. Understanding how the desialylation of the receptor microenvironment can re-tune TKI sensitivity is of paramount therapeutic importance.

In this study, we employed an integrated TMT18-labeled multi-omics pipeline to systematically map the crosstalk between sialylation and phosphorylation in A498 cells, a widely used model for ccRCC. By leveraging enzymatic in-situ remodeling of the cell-surface glycoproteins, we demonstrate that sialic acid occupancy is a prerequisite for EGF-induced signaling. Our comparative analysis reveals that sialic acid removal targets a unique phosphoproteomic program that diverges from canonical kinase inhibition by gefitinib. Furthermore, we demonstrate that targeting the glycoprotein acts as a force multiplier, significantly enhancing the efficacy of gefitinib treatment by collapsing residual signaling hubs centered on STAT3, STAT5B, and mTOR signaling. By quantifying the stoichiometric relationship between EGFR N-glycoforms and site-specific phosphorylation, and mapping these interactions within a global functional network, we provide systems-level evidence that the sialylation status of the cell surface is a fundamental regulator of the RTK interactome. Ultimately, this work highlights the potential of glycan-targeted strategies to overcome signaling bypass and therapeutic resistance in ccRCC and other RTK-driven malignancies.

## EXPERIMENTAL SECTION

### Cell culture and treatments

The A498 cells were purchased from ATCC (HTB-44) and cultured under 37°C, 5% CO₂ in minimum essential medium (MEM, Gibco, 11095080) supplemented with 10% fetal bovine serum (FBS) and 1% Penicillin-Streptomycin (Pen/Strep). For sialidase treatment, 50 μl of α2–3,6,8,9-neuraminidase A (20,000 U/mL, New England BioLabs) from Arthrobacter ureafaciens was added to the empty cell culture media (without FBS) and the treatment were performed 2h following previously established protocols^26^. After the sialidase treatment, the cells were treated with or without 50nM Gefitinib (Selleck, S1025) for 30 mins. After the Gefitinib treatment, the cells were treated with or without 50 ng/mL human epidermal growth factor (EGF, Sigma, E9644) for 15 min. After the EGF treatment, cells were washed with cold PBS on ice and harvested. For each condition, 1 million cells were obtained.

### Sample processing for protein extraction and tryptic digestion

Cellular proteins were extracted using a lysis buffer composed of 8 M urea, 75 mM NaCl, 50 mM Tris (pH 8.0), and 1 mM EDTA. To ensure protein and PTM stability, the buffer was supplemented with protease inhibitors (2 μg/ml aprotinin, 10 μg/ml leupeptin, and 1 mM PMSF), phosphatase inhibitors (10 mM NaF; 1:100 PIC2 and PIC3), and 20 μM of the O-GlcNAcase inhibitor PUGNAc. Cell pellets were processed on ice by adding 100 μl of chilled lysis buffer per pellet. Lysis was achieved through four cycles of 20-second high-speed vortexing followed by a 15-minute incubation at 4 °C. The resulting mixture was centrifuged at 20,000 g for 10 minutes at 4 °C to remove insoluble debris, and the protein-rich supernatant was collected. Protein concentrations were determined via a BCA assay (Pierce). To ensure accuracy, samples, lysis buffer controls, and water blanks were diluted 1:20 and analyzed in triplicate against a BSA standard curve. Following a 30-minute incubation at 37 °C, absorbance was recorded at 562 nm. Based on these measurements, protein concentrations were standardized to 4 μg/μl using lysis buffer, and 50 μg aliquots were prepared for downstream analysis. Protein samples underwent reduction with 6 mM dithiothreitol (DTT) for 1 hour at 37 °C, followed by alkylation using 12 mM iodoacetamide (IAA) for 45 minutes at 25 °C in a light-restricted environment. To facilitate efficient enzymatic digestion, the urea concentration was reduced to less than 2 M by diluting the samples 1:4 with 50 mM Tris-HCl (pH 8.0). Initial proteolysis was performed using LysC (Wako Chemicals) at a 1:50 enzyme-to-substrate ratio for 2 hours at 25 °C. This was followed by an overnight (∼14 hours) digestion at 25 °C using sequencing-grade modified trypsin (Promega) at the same 1:50 ratio. Post-digestion, the reaction was quenched and acidified to pH 3 using 50% formic acid (final concentration of 1% FA). The volume was then adjusted to 1 ml with 0.1% FA to promote the precipitation of any remaining undigested proteins. These precipitates were removed via centrifugation at 20,000 g for 15 minutes. The resulting tryptic peptides in the supernatant were desalted using reversed-phase C18 SPE columns (SepPak, Waters) and subsequently concentrated to dryness in a Speed-Vac.

### TMT labeling of peptides

Peptide labeling was conducted using 18-plex Tandem Mass Tag (TMT18) reagents (Thermo Fisher Scientific) to enable multiplexed quantification. For each experimental channel, 25 μg of peptides—quantified via a peptide-level BCA assay—were reconstituted in 10 μl of 100 mM HEPES (pH 8.5). TMT reagents were prepared by dissolving 5 mg of anhydrous reagent in 500 μl of anhydrous acetonitrile. Following a 5-minute equilibration period and brief vortexing, 5 μl of the TMT reagent (equivalent to 50 μg) was added to each peptide sample. This maintained a consistent 1:2 peptide-to-TMT reagent weight ratio (w/w). The labeling reaction was performed at room temperature for 1.5 hours with constant shaking at 1,000 rpm. To terminate the reaction and remove unreacted TMT tags, the samples were quenched with 2 μl of 5% hydroxylamine for 15 minutes at room temperature under continued agitation. Subsequently, the 18 labeled peptide channels were pooled into a single multiplexed sample. The combined mixture was evaporated to dryness and then reconstituted in 1 ml of 3% acetonitrile (ACN) containing 0.1% formic acid (FA). The solution was verified to be at pH 3 using indicator paper, with additional formic acid added if required. The final multiplexed sample was desalted using SepPak C18 columns to ensure purity for downstream fractionation.

### Peptide fractionation

The dried, TMT-labeled peptides were reconstituted in 900 μl of loading buffer containing 20 mM ammonium formate (NH_4_HCO_3_, pH 10) and 2% ACN. Off-line fractionation was performed using an Agilent 1200 Series HPLC system equipped with a Zorbax 300 Å Extend-C18 column (4.6 mm x 250 mm, 3.5 μm particle size; Agilent). Separation was carried out at a flow rate of 1 ml/min using a basic reversed-phase liquid chromatography (bHPLC) method. The mobile phases consisted of Solvent A (2% ACN, 5 mM NH_4_HCO_3_, pH 10) and a non-linear gradient of Solvent B (90% ACN, 5 mM NH_4_HCO_3_, pH 10) as previously described^21^. Following fractionation, the samples were bifurcated for parallel proteomic streams. A 5% aliquot of the total sample was reserved for global protein expression profiling. These fractions were evaporated to dryness and subsequently resuspended in 0.1% FA for LC-MS/MS analysis. The remaining 95% of the sample was dried down and dedicated to the enrichment of phosphopeptides for deep phosphoproteomic profiling.

### Enrichment of phosphopeptides by Fe-IMAC

The remaining 95% of the pooled peptide sample was concatenated into fewer fractions and subjected to phosphopeptide enrichment utilizing immobilized metal affinity chromatography (IMAC), following previously established protocols^27^. After enrichment, the resulting phosphopeptides were evaporated to dryness and subsequently reconstituted in 0.1% FA for LC-MS/MS analysis..

### Enrichment of glycopeptides by MAX enrichment

Following the partition of samples for phosphoproteomics, intact N-glycopeptides were selectively isolated from the remaining peptide pool. This enrichment was performed using MAX chromatography, adhering to previously established and validated protocols^28,29^. The enriched glycopeptide fractions were subsequently evaporated to dryness via vacuum centrifugation and reconstituted in 0.1% FA for high-resolution LC-MS/MS analysis..

### LC-MS/MS analysis

All mass spectrometry data were acquired using an Orbitrap Fusion Ascend Tribrid mass spectrometer (Thermo Scientific) coupled with an EvoSep One liquid chromatography system. TMT-labeled peptides from global proteome, phosphoproteome, and glycoproteome fractions were separated on a PepSep C18 column (15 cm × 150 µm, 1.5 µm particle size) and analyzed in data-dependent acquisition (DDA) mode. For global and phosphoproteomic runs, MS1 spectra were acquired at 60,000 resolution over a scan range of 350–1800 m/z, with an automatic gain control (AGC) target of 4×10⁵, a maximum injection time of 50 ms, and a charge state selection of 2–6. The DDA cycle time was set to 2 s, utilizing a dynamic exclusion period of 45 s (±10 ppm). MS2 spectra were recorded at 50,000 resolution following higher-energy collisional dissociation (HCD) at a normalized collision energy (NCE) of 34%. Precursor isolation was performed with a 0.7 m/z window, an AGC target of 2×10⁵, and a 100-ms maximum injection time. Initial glycoproteomic datasets were acquired using MS1 scans at 60,000 resolution (500–2000 m/z; AGC 5×10⁵; 50 ms) with identical cycle and exclusion settings to the proteomic runs. MS2 spectra were collected at 50,000 resolution (HCD 35%; 0.7 m/z isolation; AGC 1×10⁵; 100-ms injection). Additional glycoproteomic acquisitions were performed using an electrospray source voltage of 1.8 kV and a transfer-tube temperature of 300 °C. For these runs, MS1 resolution was increased to 120,000 (350–2000 m/z; AGC 5×10⁵), and MS2 fragmentation was executed using a stepped HCD energy (25%, 35%, and 45%). MS2 scans were recorded at 30,000 resolution with an AGC target of 3×10⁵, a 64-ms injection time, and charge-state selection optimized for glycopeptides (2–8), maintaining a 45-s dynamic exclusion window.^29^.

### Identification and quantification of phosphopeptides, glycopeptides, and proteins

Mass spectrometry raw files were processed using the MS-PyCloud software suite. Peptides were searched against the UniProtKB human reference database (version 2024.02.29). Search parameters utilized trypsin/P specificity, allowing for up to three missed cleavages for the global proteome and four for PTM-enriched datasets. Fixed modifications included carbamidomethylation of cysteine, while variable modifications included methionine oxidation, protein N-terminal acetylation, and—for phosphoproteomic data—phosphorylation of serine, threonine, and tyrosine (pSer/pThr/pTyr). The search criteria restricted peptide length to >=6 amino acids with a maximum of five modifications per peptide. Identification quality was ensured using an Andromeda score threshold of >=40 and a delta score of >=6. False discovery rates (FDR) were controlled at 1% at the peptide-spectrum match (PSM) level using a target-decoy approach. For PTM datasets, the protein-group FDR was set to 100% to strictly maintain a 1% peptide-level FDR without protein-level filtering. For global proteome datasets, a standard 1% protein-level FDR was applied. Automated correction for TMT isotope impurities was performed using manufacturer-provided specifications, and downstream analysis was conducted using the resulting evidence (PTM) or protein-group (global) tables. For glycoproteomic identification, .RAW files were first converted to .mzML format using ProteoWizard (version 3.0) with the “peakPicking” filter enabled. Database searching was performed using GPQuest (version 3.0) on a high-performance computing system (64 vCPUs, 1 TB RAM)^30^. Spectra were searched against the UniProtKB human reference list (2024.02.29) integrated with a comprehensive human N-glycan database. Search parameters included a precursor mass tolerance of 10 ppm and a fragment mass tolerance of 20 ppm. Trypsin was specified as the protease, allowing for up to two missed cleavages. Eligible peptides were restricted to lengths of 7–50 amino acids and precursor charge states of +2 to +8. Identification relied on the matching of b- and y-type ions, requiring a minimum of six matched peaks per peptide. Fixed and variable modifications were set to carbamidomethylation (+57.0215 Da) and methionine oxidation (+15.9949 Da), respectively. All glycopeptide identifications were filtered to achieve a 1% PSM-level FDR.

### Data analysis and visualization

Data analysis was performed in R using a unified workflow for the global proteome, phosphoproteome, and N-glycoproteome. Reporter-ion abundance matrices were analyzed on the log_2_ scale. Contaminant entries and features with non-positive abundance values were removed. Features were retained if they were quantified in at least two of three biological replicates in at least one treatment condition. Missing values were not imputed; statistical models were fit using the available quantitative measurements, and figure panels requiring direct feature-wise comparison across samples used complete-case feature sets. Feature identifiers were standardized before statistical analysis. Protein-level features were annotated by gene symbols. Phosphosite labels were generated from phosphopeptide sequence, modification annotation, and phosphosite index information to assign residue-level labels where localization was supported; localized phosphosites were preferentially used for figure labels and site-resolved summaries. N-glycopeptides were tracked as intact glycopeptide features, defined by gene, peptide sequence, and glycan composition. Glycan compositions were parsed into HexNAc, hexose, fucose, sialic acid, and NeuGc counts, enabling analyses stratified by sialylation and fucosylation state.

Differential abundance was tested separately for each omics layer with linear models and empirical Bayes moderation using limma^31^. A six-level treatment design fitted without an intercept, and treatment contrasts were estimated for EGF stimulation, sialidase alone, sialidase added during EGF stimulation, gefitinib added during EGF stimulation, and combined sialidase/gefitinib treatment during EGF stimulation. Empirical Bayes moderation used intensity-dependent variance modeling. The *p* values were adjusted independently for each omics layer and contrast using the Benjamini-Hochberg procedure. Unless otherwise stated, significant features were defined by adjusted *p* < 0.05 and absolute log_2_ fold change > 0.58, corresponding to an approximately 1.5-fold change.

Phosphoproteomic analyses focused on EGF-responsive signaling where appropriate. EGF-responsive phosphosites were defined as sites significant after EGF stimulation relative to untreated control, and this set was used as the background for analyses of sialidase and gefitinib effects in EGF-stimulated cells. Overlap analyses were performed at phosphosite resolution; overlap summaries used set counts and, where shown, Jaccard similarity and hypergeometric enrichment testing. For trajectory and heatmap panels, phosphosite abundances were row-wise z-scored unless relative linear abundance was explicitly shown.

Kinase activity was inferred from phosphosite-level differential results using kinase-substrate enrichment analysis with curated PhosphoSitePlus substrate annotations as implemented in KSEAapp^32,33^. Phosphosite log_2_ fold changes were converted to linear fold changes for KSEA input, and kinase-level z-scores and nominal enrichment *p* values were used to summarize inferred activity changes.

Pathway enrichment was performed using DAVID Bioinformatics Resources through the web portal^34,35^. Gene lists derived from significant phosphosite sets were submitted for GO:BP_DIRECT over-representation analysis. For directional signaling analyses, genes were selected from phosphosites that increased after EGF stimulation and decreased after sialidase, gefitinib, or combined treatment in EGF-stimulated cells. Enrichment outputs retained DAVID *p* values, Benjamini-adjusted *p* values, fold enrichment, mapped genes, and gene counts; plotted enrichment panels report term-level significance and the number of mapped genes.

Glycoproteome analyses were conducted at the intact glycopeptide level. Paired glycoform analyses grouped features sharing the same gene, peptide sequence, and non-sialic glycan backbone, allowing abundance changes to be compared across sialic acid counts within matched glycopeptide families. Reciprocal sialylation-by-fucose and fucosylation-by-sialic-acid summaries were generated from these matched families.

Phospho-glyco integration was performed by matching significant phosphosites and glycopeptides by gene. Gene-level integration used the mean log_2_ fold change across matched phosphosites and glycopeptides per gene, whereas site-pair integration retained all phosphosite-glycopeptide pairs mapped to the same gene. Correlations shown for integration analyses were computed using Pearson correlation. EGFR trajectory analyses used complete EGFR phosphosite and glycopeptide features and displayed either row-wise z-scored condition means or relative linear abundance normalized to untreated control, as indicated in the figure.

Protein interaction analysis was performed in Cytoscape using STRING interactions at confidence score 0.4^36,37^. Network nodes were genes represented by significant glycopeptide and/or EGF-responsive phosphosite changes after sialidase addition during EGF stimulation. Node attributes summarized omics-layer membership, feature counts, adjusted *p* value ranges, and log_2_ fold-change ranges. Densely connected subnetworks were identified with MCODE^38^.

Quality control was assessed for each omics layer using principal-component analysis, sample-sample Pearson correlation, and coefficient-of-variation summaries. Principal-component analysis was performed on complete quantitative feature matrices after centering and scaling features. These quality-control outputs are reported in the Supporting Information.

## RESULTS AND DISCUSSION

### Multi-omics overview of EGF, sialidase, and gefitinib responses

To systematically investigate the crosstalk between protein sialylation and EGFR signaling, we employed a multi-omics approach to profile the global proteome, phosphoproteome, and N-glycoproteome of A498 cells under combinatorial perturbations (Figure 1A). Cells were subjected to sialidase treatment (S) to enzymatically remove the sialic acid of the cell surface glycoprotein, followed by stimulation with epidermal growth factor (EGF, E) and/or inhibition by the EGFR-specific tyrosine kinase inhibitor (TKI) gefitinib (G). This factorial experimental design yielded six distinct functional groups: control (C), EGF stimulation (E), EGF with gefitinib (EG), sialidase treatment (S), sialidase with EGF treatment (ES), and sialidase with EGF and gefitinib treatment (ESG). Following cell lysis and protein digestion, peptides were labeled with TMT18 reagents. We utilized Immobilized Metal Affinity Chromatography (IMAC) and Mixed-mode Anion Exchange (MAX) to enrich phosphopeptides and N-glycopeptides, respectively, before performing high-resolution liquid chromatography-mass spectrometry (LC-MS) analysis.

**Figure 1.**
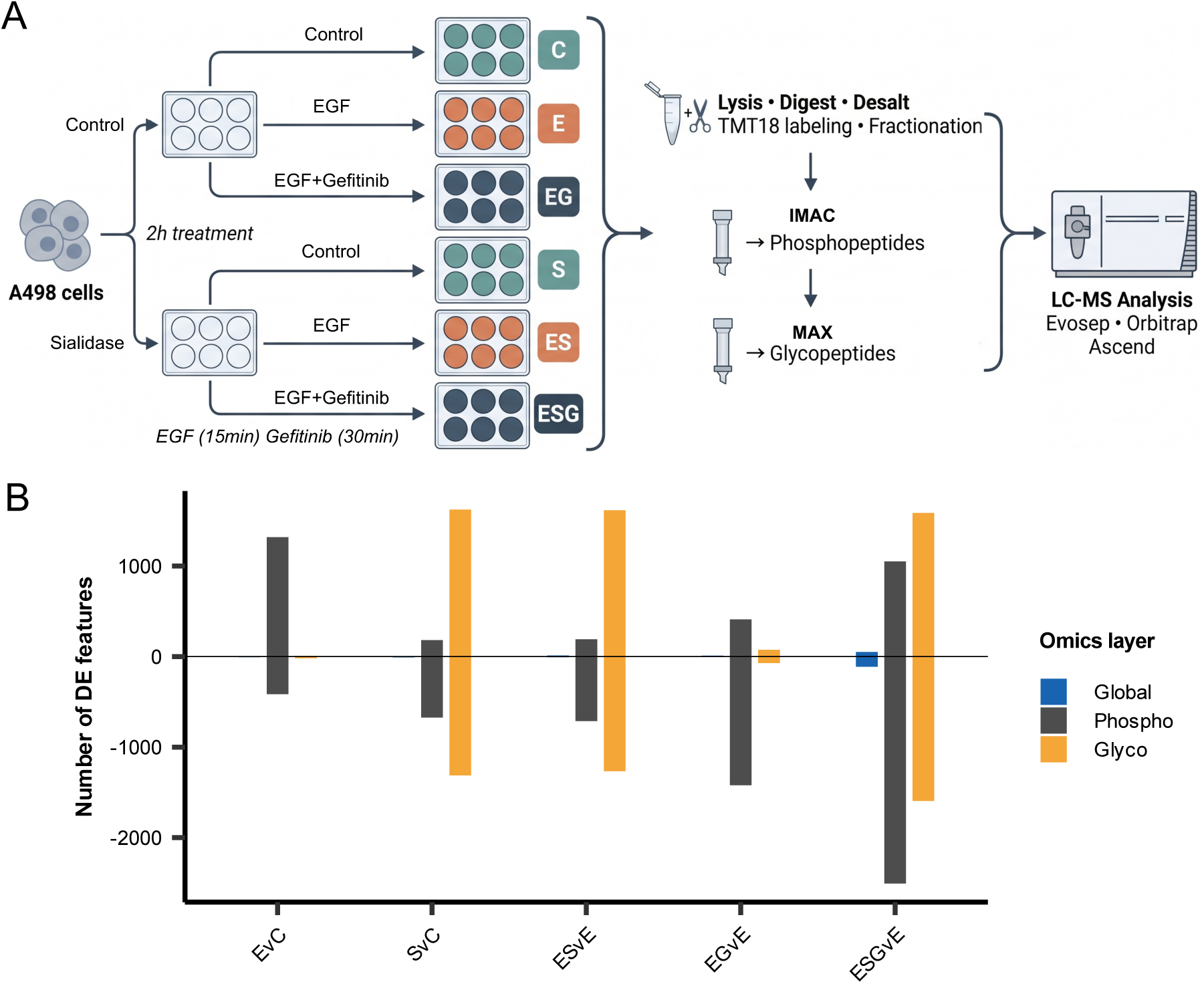
Multi-omics overview of EGF, sialidase, and gefitinib responses. (A) Experimental workflow for global proteome, phosphoproteome, and N-glycoproteome profiling. (B) Differentially expression (DE) features across treatment contrast in each omics layer. Positive and negative bars denote increased and decreased abundance, respectively; significance was defined as adjusted *p* < 0.05 and |log_2_ fold change| > 0.58.

To ensure the technical robustness of our multi-stage pipeline, we performed comprehensive quality control (QC) analysis (Supplementary Figure 1). Principal component analysis (PCA) demonstrated tight clustering of biological replicates and clear separation between treatment groups across all three omics layers (Supplementary Figure 1A). Furthermore, the coefficient of variation (CV) for all groups across the three layers remained low, indicating high technical precision (Supplementary Figure 1B). Pearson correlation analysis confirmed excellent reproducibility among triplicates, validating the quantitative accuracy of the dataset (Supplementary Figure 1C). Next we quantified differential expression (DE) features across five biologically relevant contrasts: EGF-induced activation (EvC), sialidase treatment (SvC), the impact of sialidase treatment on EGF response (ESvE), and the efficacy of Gefitinib inhibition in both native and sialidase treatment contexts (EGvE and ESGvE) (Figure 1B). Our analysis revealed that the phosphoproteome exhibited the most pronounced sensitivity to perturbation. EGF stimulation (EvC) and gefitinib inhibition (EGvE/ESGvE) triggered large-scale shifts in phosphorylation sites, highlighting the rapid signaling flux of the EGFR pathway. Sialidase treatments resulted in reorganization of the N-glycoproteome (SvC and ESvE), with thousands of glycopeptides showing significant abundance changes. This confirms successful enzymatic desialylation of the cell surface glycoprotein. In contrast to the phosphorylation and glycosylation layers, the global proteome remained stable across the 2-hour treatment window. This stability suggests that the observed changes in phosphorylation and glycosylation are a result of post-translational modifications (PTM) rather than de novo protein synthesis.

Together, these results provide a robust, multi-dimensional map of the A498 cellular landscape, setting the stage for a detailed dissection of how desialylation of the cell surface glycoprotein modulate EGFR phosphorylation dynamics and therapeutic sensitivity.

### Sialidase treatment remodels EGF-stimulated phosphorylation

To define the specific impact of the desialylation of the cell surface glycoprotein on EGF-induced signaling, we analyzed the phosphosite dynamics within the EGFR signaling (Figure 2). Initial stimulation with EGF (EvC) triggered a robust signaling response, with 1,318 upregulated and 415 downregulated phosphosites (|log_2_ FC| > 0.58, adjusted *p* < 0.05). Notably, key regulatory phosphosites such as EGFR Y1197 and S1166, as well as STAT5B Y699, were significantly upregulated, confirming the activation of canonical RTK pathways (Figure 2A). We then investigated how sialidase treatment modulates this EGF-responsive landscape (ESvE). A subset of EGF-responsive phosphosites exhibited significant sensitivity to sialidase treatment, with 33 phosphosites showing increased and 207 phosphosites showing decreased phosphorylation levels upon sialidase treatment (Figure 2B). Notably, EGFR Y1197 demonstrated significantly downregulated phosphorylation in sialidase treated cells, suggesting that the sialic acids of the cell surface glycoproteins are required for the full magnitude of EGF-stimulated signaling. Venn diagram analysis revealed an overlap between EGF-responsive and sialidase-modulated phosphosites, with 240 phosphosites being co-regulated by both perturbations (Figure 2C). To identify the upstream regulators driving these changes, we performed kinase-substrate enrichment analysis (KSEA) based on the ESvE contrast. Sialidase treatment led to a significant downregulation of several key kinases, including AURKB, MAP2K3, and CDK1, whereas a smaller subset of kinases like MAPK15 and PLK3 showed increased activity (Figure 2D). This indicates that the desialylation of the cell surface glycoprotein does not merely dampen all signaling but rather selectively reprograms specific kinase-substrate networks. A detailed examination of EGFR-centered signaling components using a heatmap across all conditions (C, E, S, ES) further highlighted the suppressive effect of sialidase treatment on key nodes (Figure 2E). We observed that EGF-induced phosphorylation of critical proteins including MAP2K2 (S226), MAPK3 (T202/Y204), SHC1 (Y349/Y350), and RAF1 (S25) were consistently diminished by sialidase treatment. Gene Ontology (GO) enrichment analysis of these phosphosites revealed that they are primarily involved in actin cytoskeleton organization, cell migration, and the MAPK cascade (Figure 2F).

**Figure 2.**
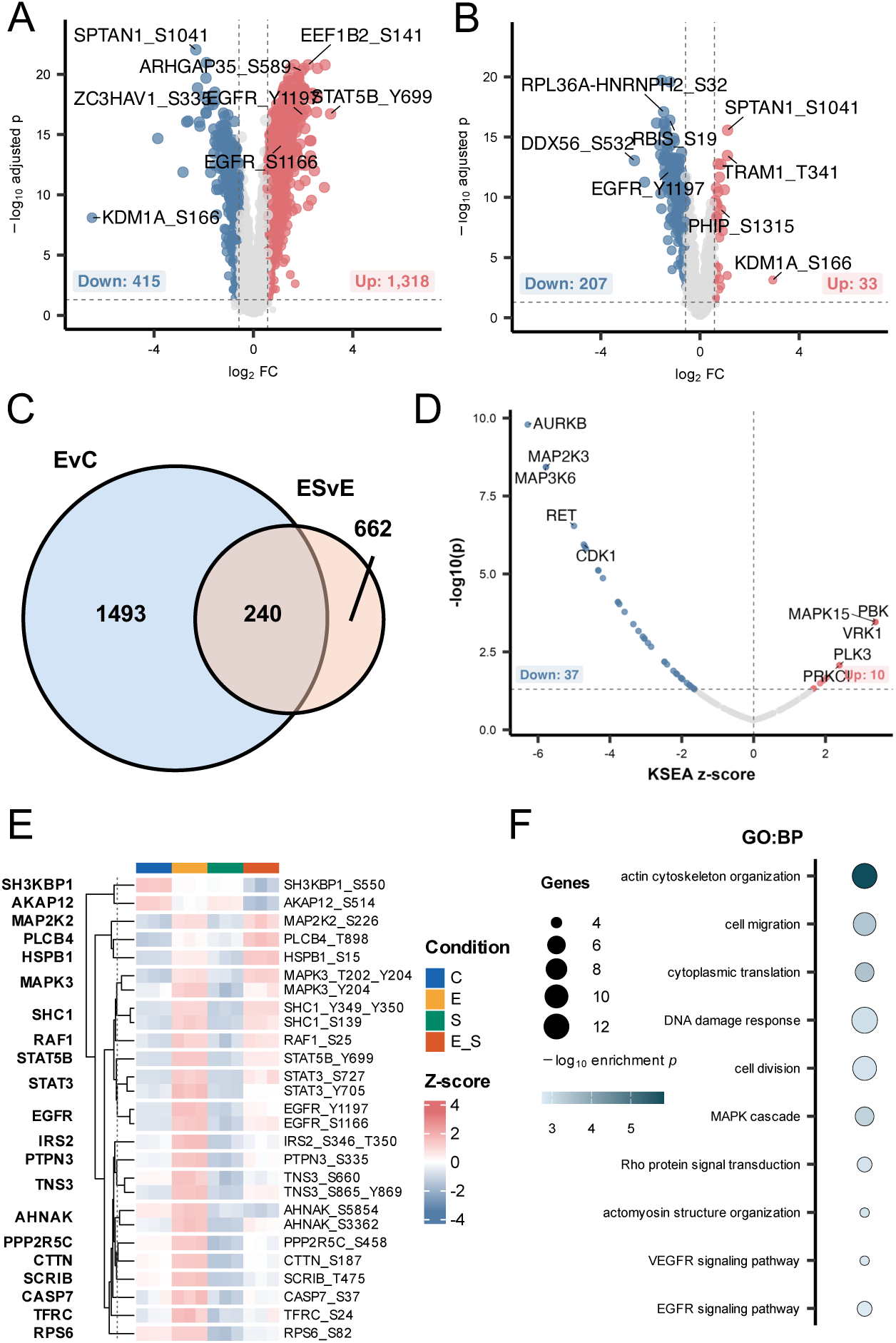
Sialidase treatment remodels EGF-stimulated phosphorylation. (A) Phosphosite changes induced by EGF. (B) Sialidase-dependent changes among EGF-responsive phosphosites. (C) Site-level overlap between EGF-responsive and sialidase-modulated phosphosites. (D) Kinase substrate enrichment after sialidase addition during EGF stimulation. (E) EGFR-centered phosphosite heatmap across control, EGF, sialidase, and combined EGF plus sialidase conditions. (F) Biological processes enriched among EGF-induced phosphosites reversed by sialidase.

In summary, these data demonstrate that the sialic acids of the cell surface glycoprotein play a crucial role in maintaining the intensity and duration of EGF-stimulated signaling, particularly within the MAPK pathway and cytoskeleton-related processes, suggesting that glycoproteome remodeling is a potent regulator of RTK-mediated cellular responses. The observed attenuation of EGFR phosphorylation at Y1197 upon desialylation is consistent with the previous findings which demonstrated that hypersialylation promotes EGFR activation and confers resistance to gefitinib^39,40^. Furthermore, the enrichment of these phosphosites in actin cytoskeleton organization (Figure 2F) suggests a potential disruption of the galectin-glycan lattice, a mechanism previously shown to regulate the surface residence and signaling duration of RTKs^41,42^.

### Sialidase and gefitinib treatments reveal shared and distinct EGF-dependent phosphosite programs

To evaluate the extent to which sialidase treatment mimics or diverges from pharmacological EGFR inhibition, we compared the phosphoproteome shifts induced by sialidase treatment (ESvE) and gefitinib treatment (EGvE) (Figure 3). Fold change analysis revealed a low global correlation, indicating that sialidase and gefitinib modulate largely distinct subsets of the phosphoproteome (Figure 3A). However, a cohort of phosphosites was co-regulated by both treatments (indicated in purple), suggesting a shared regulatory axis for a specific signaling core. A three-way Venn diagram analysis was performed to intersect EGF-responsive phosphosites (EvC) with those modulated by sialidase treatment (ESvE) and gefitinib treatment (EGvE) (Figure 3B). This analysis identified 112 phosphosites that were significantly regulated across all three conditions. Interestingly, 128 phosphosites were uniquely co-regulated by EGF and sialidase treatment, independent of gefitinib treatment, while 993 phosphosites were shared between EGF stimulation and gefitinib treatment, representing the canonical, TKI-sensitive EGFR signaling pathway. To dissect the functional implications of these distinct clusters, we utilized pathway-annotated heatmaps. Phosphosites specifically modulated by sialidase treatment (ESvE) but not significantly impacted by gefitinib treatment were enriched in processes such as actin cytoskeleton organization, cell-substrate adhesion (e.g., AHNAK, CTTN), and RNA splicing (Figure 3C). Profile plots for these phosphosites (Figure 3E) show a characteristic sialidase-sensitive trajectory where phosphorylation levels are maintained or increased by gefitinib treatment but decreased upon sialidase treatment. The 112 phosphosites shared between sialidase and gefitinib treatments (Figure 3D) were highly enriched in the MAPK cascade, RTK signaling, and cell cycle regulation. This cluster includes key regulatory nodes such as EGFR (Y1197), MAPK1/3 (T202/Y204), and STAT3 (S727). Trajectory analysis (Figure 3F) reveals that these phosphosites are EGF-inducible and similarly suppressed by both enzymatic desialylation and pharmacological TKI treatment.

**Figure 3.**
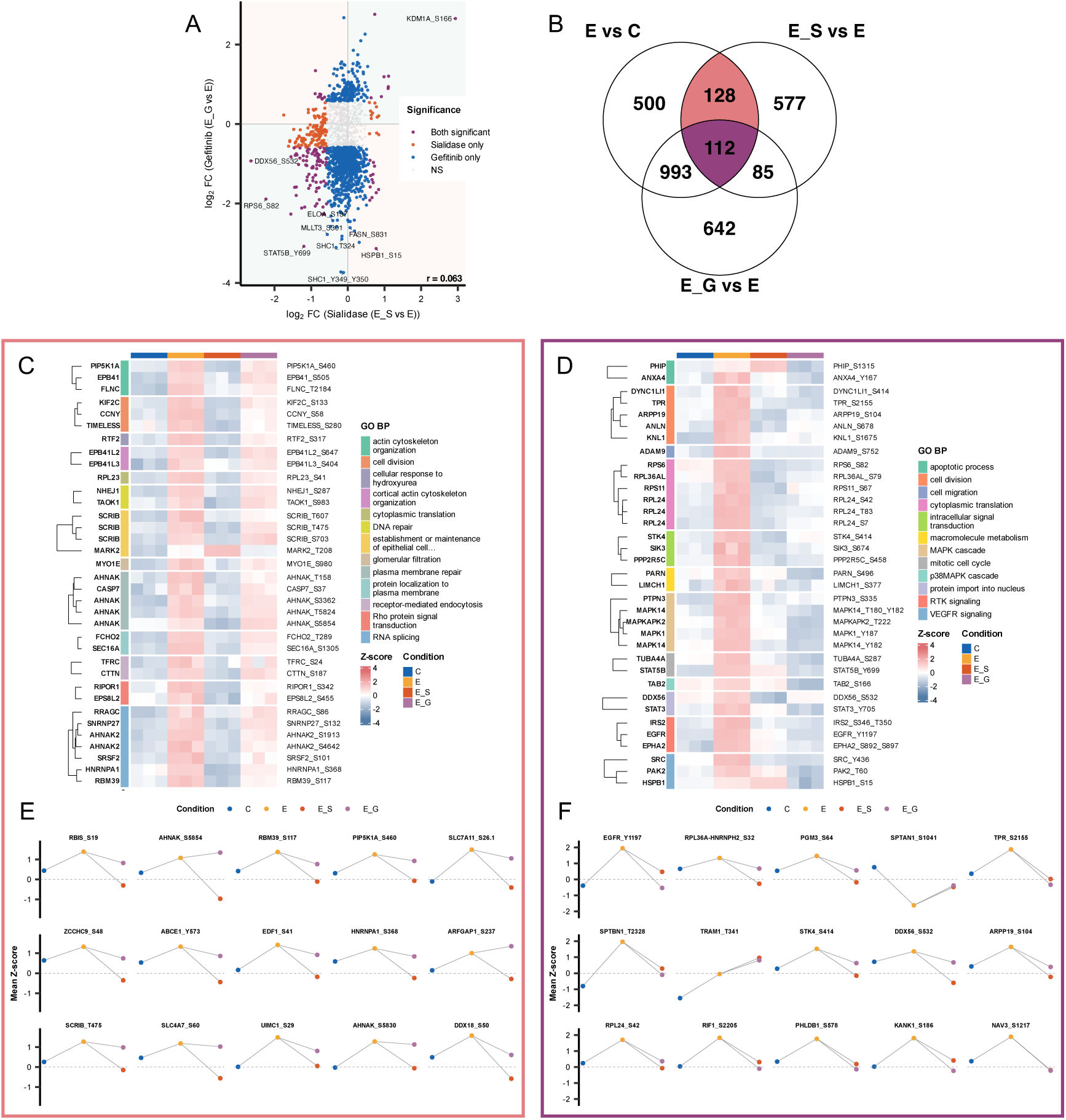
Sialidase and gefitinib treatments reveal shared and distinct EGF-dependent phosphosite programs. (A) Comparison of phosphosite log_2_ fold changes after sialidase or gefitinib treatment in EGF-stimulated cells. (B) Three-way overlap of significant phosphosites across EGF stimulation, sialidase modulation, and gefitinib inhibition. (C,D) Pathway-annotated heatmaps of sialidase-specific and shared sialidase/gefitinib-responsive phosphosites. (E,F) Representative phosphosite trajectories from the corresponding overlap regions.

The stark divergence between sialidase- and gefitinib-induced phosphoproteomic profiles underscores a fundamental decoupling of surface-glycan regulation from canonical kinase catalysis. While gefitinib provides targeted inhibition of the EGFR intracellular kinase domain, enzymatic desialylation selectively destabilizes a glycan-dependent signaling axis. This is evidenced by the attenuation of cytoskeletal and adhesion-related phosphonodes (e.g., AHNAK, CTTN) that remain refractory to gefitinib, suggesting a non-redundant regulatory mechanism. These effects likely stem from the loss of terminal sialic acids, which are essential for the structural integrity of the galectin-glycan lattice. This supramolecular scaffold maintains the spatiotemporal organization of RTKs at the plasma membrane, preventing aberrant endocytosis and off-target interactions^43^. Furthermore, the selective reprogramming rather than generalized suppression of kinase-substrate networks supports the paradigm of glycosylation-dependent signaling bias, where the glycan shield acts as a molecular gatekeeper to prioritize specific downstream effectors. Collectively, these data identify the glyco-code as a high-precision therapeutic target for modulating tumor cell migration, offering a mechanistic alternative to the systemic limitations of traditional TKIs.

### Combined sialidase and gefitinib treatment defines a distinct phosphoproteomic state

To characterize the integrated impact of sialidase treatment and tyrosine kinase inhibition, we analyzed the phosphoproteome under the combined condition of sialidase and gefitinib treatment (ESG). An UpSet plot analysis was employed to visualize the intersections of differentially abundant phosphosites across the primary contrasts (Figure 4A). The analysis revealed that the ESGvE contrast yielded the largest set of modulated phosphosites, with a substantial portion (over 1,500 sites) being unique to the combined treatment. This suggests that the interplay between sialic acid removal and TKI inhibition does not merely result in the sum of individual effects but rather activates a distinct phosphoproteomic program. A global comparison of the magnitude of phosphorylation changes (Figure 4B) demonstrated that the combined E+S+G treatment induced a more pronounced downward shift in fold changes compared to either sialidase (E+S) or gefitinib (E+G) alone. This global trend indicates a potentiated inhibitory effect on the signaling landscape when both the receptor’s extracellular glycans and its intracellular kinase activity are targeted. To identify the specific signaling nodes most affected by this combination, we examined representative phosphosites that exhibited an additive inhibitory response (Figure 4C). Critical regulatory sites, including RPS6 (S82) showed significantly greater suppression in the ESG condition compared to single treatments. This additive effect suggests that sialic acids may facilitate residual EGFR signaling that is partially resistant to gefitinib alone, and their removal further sensitizes these core nodes to inhibition. Functional enrichment analysis of the EGF responsive phosphosites suppressed by the combined treatment highlighted a broad impact on essential cellular pathways (Figure 4D). These included actin cytoskeleton organization, intracellular signal transduction, and the MAPK cascade. Interestingly, when focusing on phosphosites that were uniquely altered in the additive ESG state, we observed a specific enrichment in cell division and mitotic cycle regulation, suggesting a potent block on proliferative signaling.

**Figure 4.**
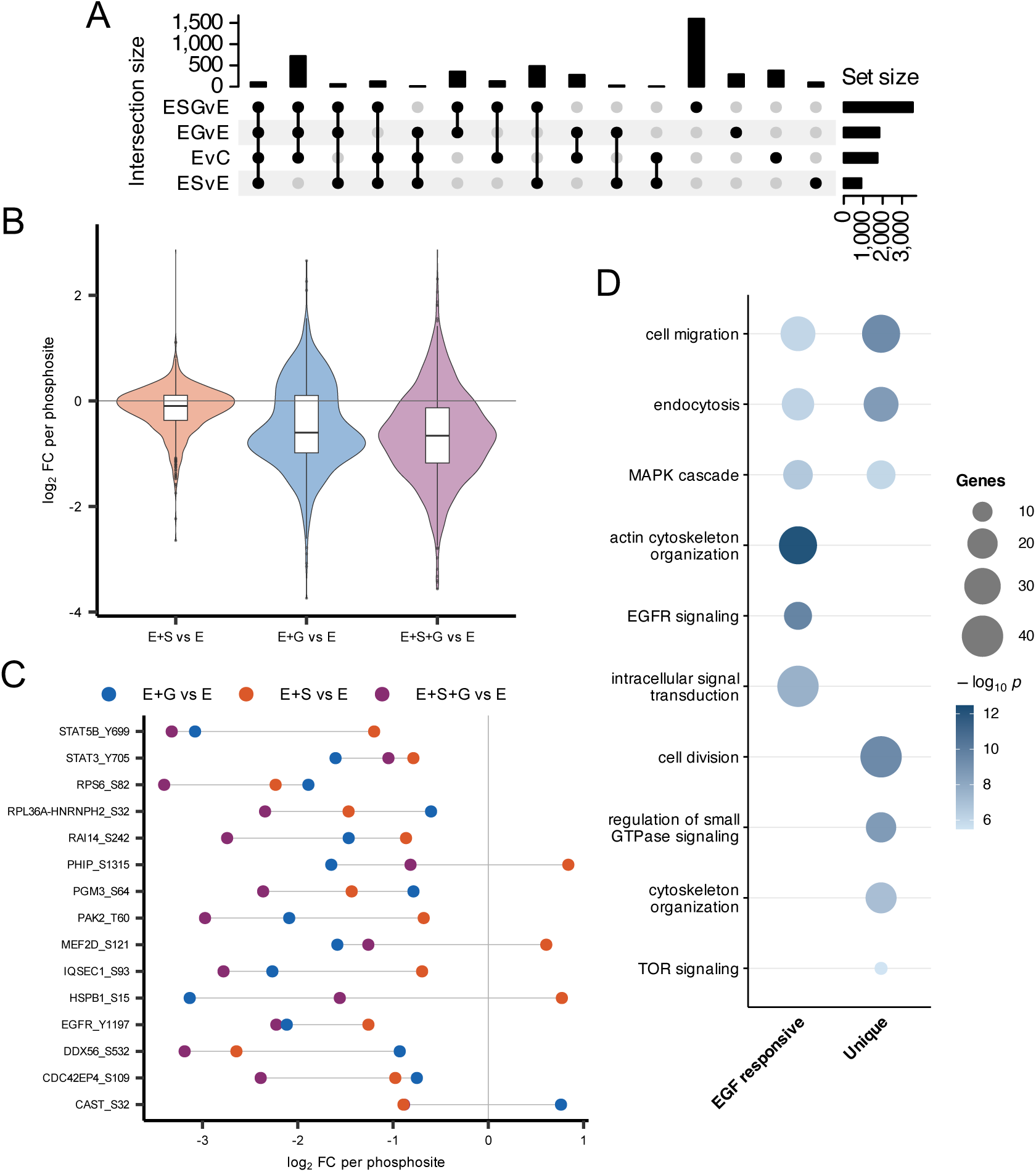
Combined sialidase and gefitinib treatment defines a distinct phosphoproteomic state. (A) UpSet analysis of significant phosphosites across EGF stimulation, sialidase, gefitinib, and combined treatment. (B) Distribution of phosphosite log_2_ fold changes for the single and combined perturbations. (C,D) Biological processes enriched among EGF-induced phosphosites suppressed by combined treatment and among phosphosites uniquely altered by combined treatment.

Collectively, these data demonstrate that sialidase treatment acts as a force multiplier for gefitinib, precipitating a more comprehensive collapse of the EGFR-driven signaling network than either agent alone. The emergence of unique functional enrichments in the combined state, particularly the potent block on mitotic cycle regulation, suggests that targeting cell surface glycoproteins may bypass compensatory signaling pathways that typically drive TKI resistance. This potentiated inhibition aligns with the evidence that hypersialylation constitutes a protective glycan shield that sustains residual RTK activity under pharmacological challenge by modulating receptor density and ligand-binding kinetics^44,45^. Specifically, the synergistic suppression of RPS6 supports the hypothesis that terminal sialic acids facilitate a bypass signaling mechanism; for instance, the sialyltransferase ST6Gal-I has been shown to mediate EGFR-TKI resistance^40^. Furthermore, the unique impact of the combined ESG treatment on cell division processes suggests that desialylation may disrupt the integrin-RTK crosstalk required for cell cycle progression. By integrating these findings, our data position enzymatic glycan stripping as a strategic necessity to overcome the intrinsic glyco-resistance of tumor cells, effectively extinguishing signaling nodes that are typically refractory to intracellular kinase inhibition alone.

### EGFR-linked phosphorylation change is coupled with Glycoproteome remodeling

To provide a mechanistic link between extracellular glycan of the cell surface glycoprotein and intracellular signaling, we analyzed the N-glycoproteome response to sialidase treatment in the context of EGF stimulation (Figure 5). Sialidase treatment induced massive shifts in the glycoproteome, with 1,615 upregulated and 1,265 downregulated glycopeptides (Figure 5A). Notably, prominent cell-surface receptors and adhesion molecules, such as CD44, LAMP2, and EGFR, exhibited significant glycan remodeling. We further characterized these changes by analyzing the sialylation state of the identified glycoforms. A ranked response plot revealed that the directionality of the glycopeptide abundance change was strictly dependent on the initial sialic acid count (Figure 5B). Highly sialylated glycoforms (S>=3) were depleted, while non-sialylated (S=0) or low-sialylated glycoforms (S=1) increased in relative abundance. This enzymatic shaving of sialic acids was consistent across different fucosylation levels (F=0 to F>=3), as shown by the stratified trajectories in Figure 5C, confirming that sialic acid removal was the primary driver of glycoproteomic variance. To determine if these glycan shifts directly coordinate with signaling changes, we performed a correlation analysis between glycopeptide and phosphopeptide abundances at both the gene and site-pair levels (Figure 5D, E). For key RTKs such as EGFR, we observed that the sialidase-induced depletion of sialic acids from N-glycopeptides was directly accompanied by a concomitant reduction in the phosphorylation levels of these specific receptors. Detailed examination of EGFR-specific trajectories (Figure 5F) provided the most compelling evidence for glyco-phospho coupling. At the phosphorylation level, EGF-induced activation of Y1197 and S1166 was significantly attenuated by prior sialic acid removal. Correspondingly, at the glycosylation level, we observed sialidase treatment depleted complex, sialylated glycoforms at residues N413, N444, and N603, while leading to a compensatory increase in simpler, desialylated structures.

**Figure 5.**
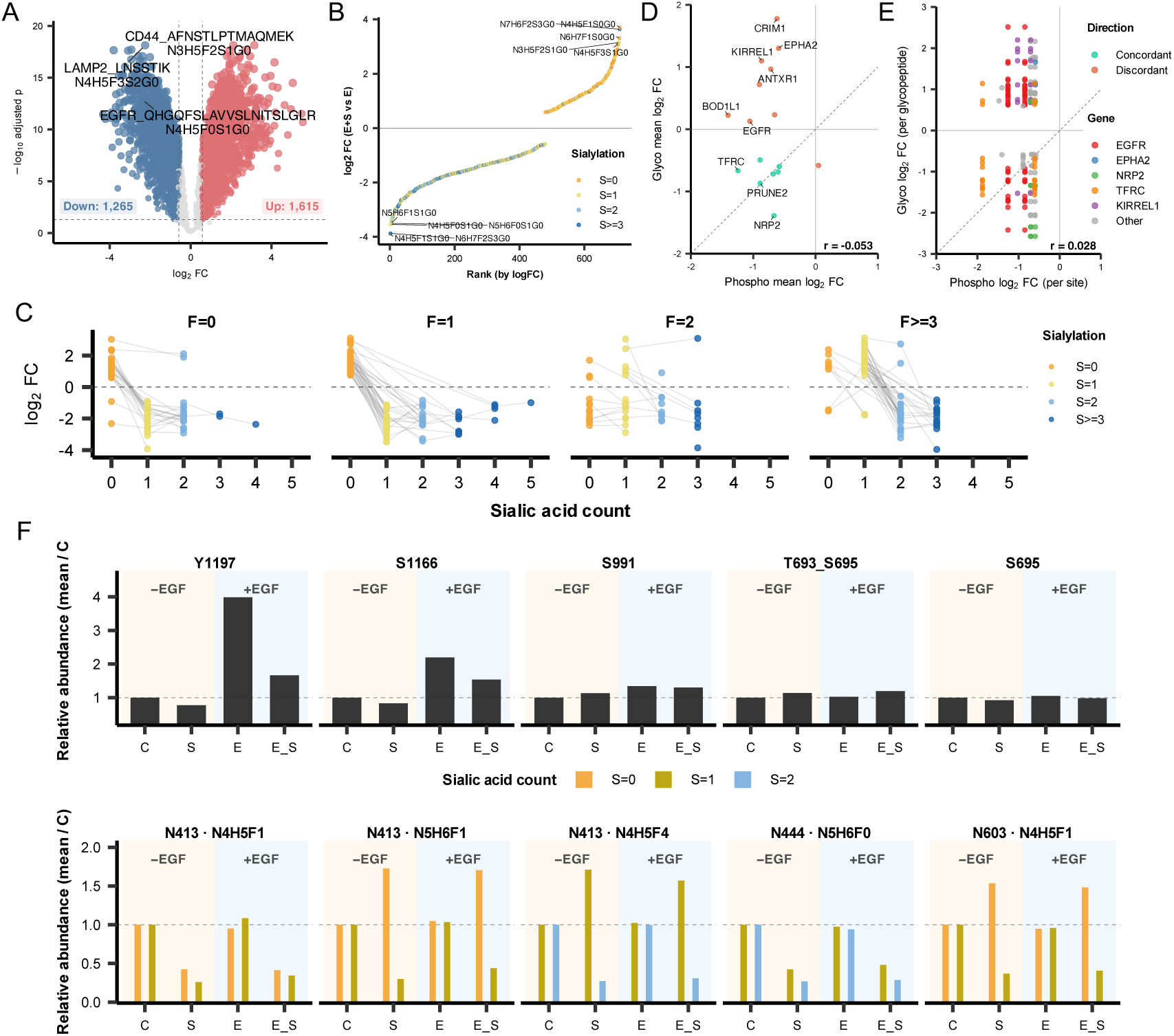
EGFR-linked phosphorylation change is coupled with Glycoproteome remodeling. (A) Glycopeptide abundance changes after sialidase addition during EGF stimulation. (B) Ranked paired-glycoform response plot colored by sialic acid count. (C) Sialylation-dependent glycopeptide changes stratified by fucose count. (D,E) Gene-level and site-pair comparisons of phosphorylation and glycosylation responses. (F) Relative-abundance trajectories for selected EGFR phosphosites and EGFR N-glycoforms.

Collectively, these findings demonstrate a direct mechanistic coupling between the extracellular glycan architecture and intracellular phosphorylation dynamics of EGFR. The systematic depletion of high-order sialylated glycoforms and the concomitant reduction in Y1197/S1166 phosphorylation (Figure 5F) suggest that terminal sialic acids are indispensable for sustaining the receptor in a signaling-competent state. This stoichiometric relationship supports the hypothesis that the sialic acid cap on N-glycans modulates the conformational equilibrium of the EGFR extracellular domain. The loss of these negatively charged moieties may increase the energetic barrier for ligand-induced dimerization or alter the receptor’s orientation within the plasma membrane^46,47^. Furthermore, the observation that sialic acid depletion rather than the total loss of the glycan tree is sufficient to attenuate signaling aligns with the concept of charge-dependent receptor gating, where the electrostatic environment created by sialylation prevents the receptor from adopting an autoinhibited or tethered conformation^48,49^. These data provide a site-specific blueprint of how glycan remodeling serves as a molecular switch for RTK activity, positioning terminal sialylation as a tunable rheostat for growth factor responsiveness in oncology.

### Integrated glyco-phospho protein interaction network

To visualize the global connectivity between the N-glycoproteome and the phosphoproteome, we constructed a protein-protein interaction (PPI) network based on genes that exhibited significant sialidase-modulated glycopeptide and/or EGF-responsive phosphosite changes (Figure 6). Using the STRING database and MCODE clustering analysis, we identified a highly interconnected network containing 800 significant features across 614 glycopeptides and 167 phosphosites, with 19 genes exhibiting significant alterations in both omics layers simultaneously. Our topological analysis identified four primary functional modules that define the desialylation-sensitive landscape of A498 cells. A core signaling hub was identified involving EGFR, AXL, ERBB2, and ITGB1. This module highlights a dense cluster of receptor tyrosine kinases and cell-surface adhesion proteins. The presence of significant glycan remodeling on these receptors, coupled with suppressed phosphorylation nodes, suggests that sialylation is critical for maintaining the structural and functional integrity of these signaling complexes at the plasma membrane. A second module centered around CD44, MET, and multiple integrin subunits (e.g., ITGA4, ITGA5, ITGB3). This suggests that desialylation may disrupt the crosstalk between growth factor receptors (MET) and adhesion-mediated signaling pathways, potentially impacting cell migration and immune evasion mechanisms. A distinct cluster was composed of ribosomal proteins (e.g., RPL23, RPS11, RPS4X) and elongation factors. The modulation of these intracellular components, primarily through phosphorylation, indicates that surface glycan remodeling triggers a retrograde signal that reaches the cellular protein synthesis apparatus, possibly as a long-term adaptive response to altered membrane signaling. A dedicated module involving ICAM1, VCAM1, and CD47 were enriched. These proteins are vital for cell-cell interactions and immune signaling. The significant changes in their glycan profiles upon sialidase treatment point toward a role for sialylation in masking or modulating the accessibility of these receptors to their cognate ligands or neighboring cells.

**Figure 6.**
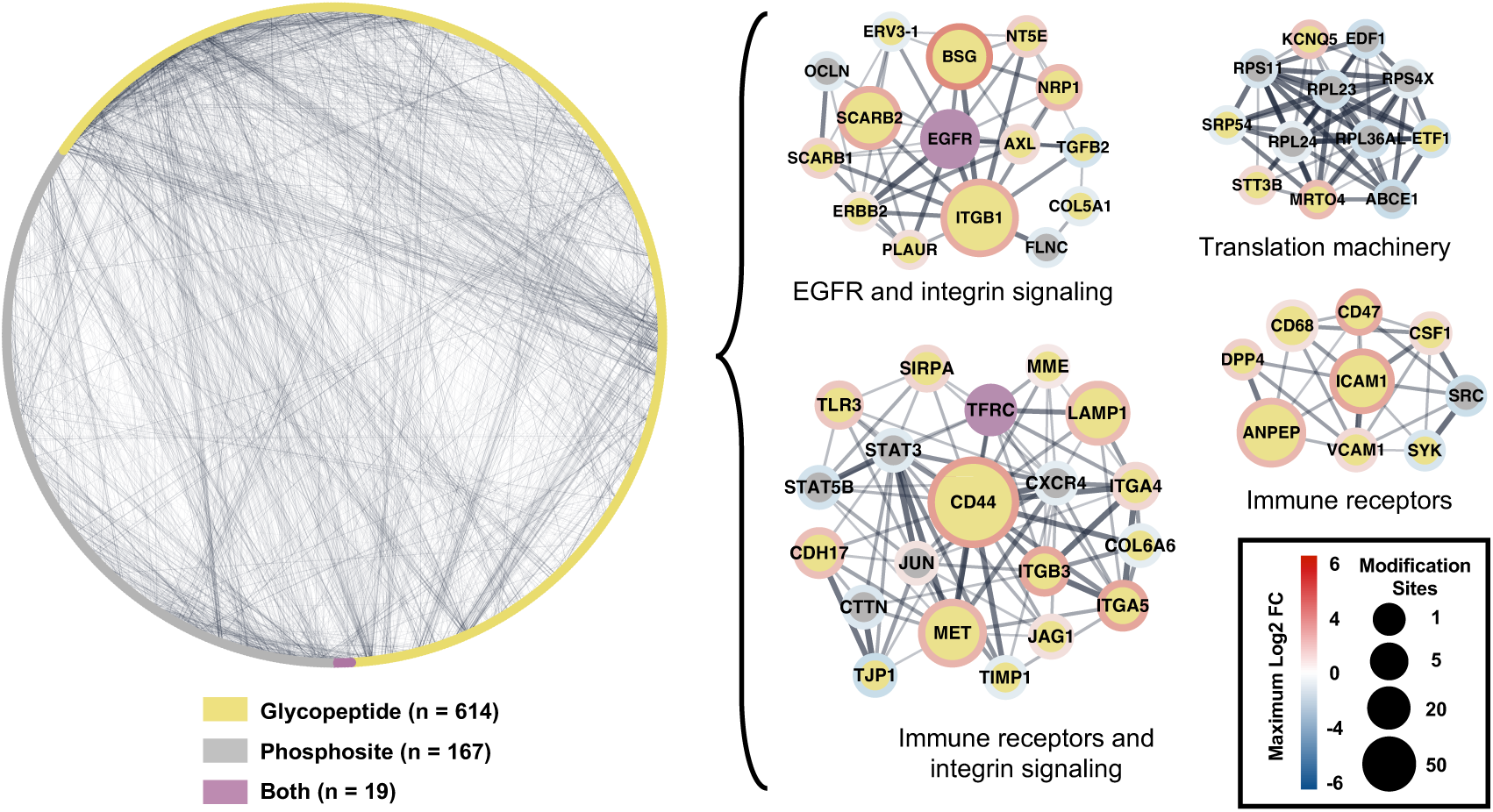
Integrated glyco-phospho protein interaction network. STRING interactions at confidence score 0.4 were used to build a Cytoscape network from genes with significant glycopeptide and/or EGF-responsive phosphosite changes after sialidase addition during EGF stimulation. Node attributes encode omics layer, feature counts, adjusted *p* ranges, and log_2_ fold-change ranges; MCODE was used to identify densely connected modules.

In summary, the integrated network demonstrates that sialic acid removal does not impact the cell in an isolated manner; rather, it triggers a coordinated systemic response across diverse functional modules. The identification of four distinct yet interconnected clusters ranging from RTK/Integrin hubs to the intracellular translation machinery establishes the cell-surface glycome as a master regulator of the cellular interaction landscape. This global connectivity supports the emerging concept of the glyco-phospho-interactome, where surface sialylation provides a necessary scaffold that coordinates signaling across multiple omics layers^50^. The modulation of ribosomal protein phosphorylation in response to surface desialylation is particularly striking, suggesting a retrograde signaling axis that couples membrane glycan status to the cell’s biosynthetic capacity. Such long-range omics crosstalk may be mediated by the MAPK/RSK pathway, which acts as a bridge between receptor-level perturbations and the regulation of the protein synthesis apparatus^51^. Furthermore, the significant glycan remodeling of immune-modulatory nodes like CD47 and ICAM1 suggests that the sialylation state may function as a glycan checkpoint, determining the receptor’s accessibility and its subsequent ability to recruit intracellular signaling effectors^52,53^. Collectively, these findings provide a topological blueprint of the desialylation-sensitive landscape, positioning surface glycosylation as the fundamental architect of coordinated cellular responses in oncogenic signaling.

## CONCLUSION

In conclusion, this study provides a multi-dimensional, high-resolution atlas of the functional interplay between cell-surface glycosylation and intracellular signaling dynamics. By integrating global proteomic, phosphoproteomic, and N-glycoproteomic profiling, we demonstrate that while the total proteome remains largely refractory to acute sialic acid perturbation, the cell-surface sialylation status acts as a critical rheostat, intrinsically coupling extracellular glycan architecture to the intensity and specificity of downstream signal transduction (Figure 1).

Our findings reveal that enzymatic desialylation significantly attenuates the canonical EGF response by selectively suppressing key regulatory phosphonodes within the MAPK cascade and actin cytoskeleton organization pathways. This re-tuning effect highlights a mechanistic decoupling of glycan-mediated regulation from direct kinase catalysis. Notably, the phenotypic divergence between sialidase treatment and pharmacological EGFR inhibition (Gefitinib) underscores that surface glycans regulate a distinct suite of cytoskeletal and structural proteins—such as AHNAK and CTTN—that are largely insensitive to TKI treatment alone (Figures 2 and 3).

Furthermore, we establish a potent synergistic axis between enzymatic desialylation and TKI inhibition. The combined treatment (ESG) precipitates a comprehensive collapse of the EGFR-driven network, driving cells into a unique phosphoproteomic state characterized by an additive suppression of critical regulators (e.g., RPS6) and a unique block on mitotic progression (Figure 4). These data position the glycan shield as a facilitatory layer that sustains residual signaling under therapeutic pressure, thereby identifying it as a key driver of intrinsic glyco-resistance.

At the molecular level, we provide site-specific evidence of glyco-phospho coupling on EGFR. The depletion of complex, sialylated N-glycans at specific residues (e.g., N413, N444, and N603) directly correlates with the dampening of activation-linked phosphosites Y1197 (Figure 5). This interaction is topologically reinforced within a global protein interaction network, where sialylation maintains the functional connectivity of hub modules involving EGFR, MET, CD44, and integrins (Figure 6).

Ultimately, this work establishes that terminal sialylation is a prerequisite for robust RTK signaling. By deconstructing the crosstalk between these two major post-translational modifications, we identify the cell-surface glyco-code as a high-precision therapeutic target, offering a strategic avenue to enhance TKI efficacy and overcome signaling bypass mechanisms in oncology.

## SUPPORTING INFORMATION

Supporting Information: Figure S1. Multi-omics quality-control summary: principal-component analysis, sample-sample correlations, and coefficient-of-variation distributions for global proteome, phosphoproteome, and glycoproteome datasets. Figure S2. Fucosylation-resolved glycopeptide responses to sialidase within matched glycopeptide families stratified by sialic acid count. Table S1. Differential-response counts for phosphosite, glycopeptide and global protein features across the manuscript treatment contrasts. Table S2. Phosphoproteomic data supporting the sialidase modulation of EGF-stimulated phosphorylation, including volcano-plot values, phosphosite overlap counts, kinase substrate enrichment results, EGFR-network heatmap values, GO Biological Process enrichment, and directional gene and phosphosite lists. Table S3. Phosphosite comparison data for sialidase and gefitinib responses in EGF-stimulated cells, including double-logFC values, three-way overlap counts, pathway heatmap matrices, representative phosphosite profiles, and Venn-region phosphosite lists. Table S4. Combined sialidase/gefitinib phosphoproteomic response data, including UpSet phosphosite membership, phosphosite log_2_ fold-change distributions, GO Biological Process enrichment, and combined-treatment-unique phosphosite lists. Table S5. Glycoproteome and phospho-glyco integration data, including glycopeptide volcano values, paired glycoform response values, sialylation-by-fucose summaries, gene-level and site-pair phospho-glyco comparisons, and EGFR relative-abundance trajectories. Table S6. STRING/Cytoscape network support tables, including glyco-phospho node attributes and glycopeptide heatmap values for genes represented in the integrated protein-interaction analysis. Table S7. Quality-control data for the supporting multi-omics QC figure, including PCA coordinates, sample correlation matrices, and coefficient-of-variation summaries for global proteome, phosphoproteome, and glycoproteome datasets. Table S8. Fucosylation-by-sialic-acid glycopeptide family data supporting the reciprocal glycan-composition analysis in Figure S2.

## AUTHOR INFORMATION

### DATA ACCESSIBILITY

The mass spectrometry proteomics data have been deposited to the ProteomeXchange Consortium via the PRIDE^54^ partner repository with the data set identifier PXD078365. Reviewer could log in to the PRIDE website using the following details: Project accession: PXD078365, Token: lyVbeSqKcT4b

## ACKNOWLEDGMENTS

This work was supported by National Institutes of Health, National Cancer Institute, the Clinical Proteomic Tumor Analysis Consortium (CPTAC, U24CA271079), the Early Detection Research Network (EDRN, U2CCA271895), Pancreatic Cancer Detection Consortium (PCDC, U01CA274514). This manuscript is the result of funding in whole or in part by the National Institutes of Health (NIH). It is subject to the NIH Public Access Policy. Through acceptance of this federal funding, NIH has been given a right to make this manuscript publicly available in PubMed Central upon the Official Date of Publication, as defined by NIH.

